# Synthetic chromosome fusion: effects on genome structure and function

**DOI:** 10.1101/381137

**Authors:** Jingchuan Luo, Luis A. Vale-Silva, Adhithi R. Raghavan, Guillaume Mercy, Jonna Heldrich, Xiaoji Sun, Mingyu Li, Weimin Zhang, Neta Agmon, Kun Yang, Jitong Cai, Giovanni Stracquadanio, Agnès Thierry, Yu Zhao, Camila Coelho, Stephanie Lauer, Ju Young Ahn, Greg Adoff, Andrew D’Avino, Henri Berger, Yi Chen, Michael Chickering, Oren Fishman, Rebeca Vergara Greeno, Sangmin Kim, Sunghan Kim, Hong Seo Lim, Jay Im, Lauren Meyer, Allison Moyer, Surekha Annadanam, Natalie A. Murphy, Peter Natov, Maisa Nimer, Arthur Radley, Arushi Tripathy, Tony Wang, Nick Wilkerson, Tony Zheng, Vivian Zhou, Karen Zeller, David B. Kaback, Joel S. Bader, Leslie A. Mitchell, Julien Mozziconacci, Romain Koszul, Andreas Hochwagen, Jef D. Boeke

**Affiliations:** Institute for Systems Genetics and Department of Biochemistry and Molecular Pharmacology, NYU Langone Health, New York, NY 10016, USA; Biochemistry, Cellular and Molecular Biology Graduate program, Johns Hopkins University School of Medicine, Baltimore, MD 21205, USA; Institut Pasteur, CNRS UMR3525, Université de Paris, Unité Régulation Spatiale des Génomes, F-75015 Paris, France; Department of Biomedical Engineering, Tandon School of Engineering, Brooklyn NY 11201; Department of Biology, New York University, New York, NY 10003, USA; High Throughput Biology Center, Johns Hopkins University School of Medicine, Baltimore, MD 21205, USA; Department of Biomedical Engineering and Institute of Genetic Medicine, Whiting School of Engineering, JHU, Baltimore, MD 21218, USA; School of Biological Sciences, The University of Edinburgh, Edinburgh, UK; Build A Genome Class, Krieger School of Arts and Sciences and Whiting School of Engineering, Johns Hopkins University, Baltimore, MD 21218, USA; Structure and instability of Genomes Lab, UMR 7196, Muséum National d’Histoire Naturelle (MNHN), 75005 Paris; Department of Microbiology and Molecular Genetics, Rutgers New Jersey Medical School, International Center for Public Health, Newark, New Jersey 07101-1709; Collège Doctoral, Sorbonne Université, F-75005 Paris, France

## Abstract

As part of the Synthetic Yeast 2.0 (Sc2.0) project, we designed and synthesized synthetic chromosome I. The total length of synI is ∼21.4% shorter than wild-type chromosome I, the smallest chromosome in *Saccharomyces cerevisiae*. SynI was designed for attachment to another synthetic chromosome due to concerns of potential instability and karyotype imbalance. We used a variation of a previously developed, robust CRISPR-Cas9 method to fuse chromosome I to other chromosome arms of varying length: chrIXR (84kb), chrIIIR (202kb) and chrIVR (1Mb). All fusion chromosome strains grew like wild-type so we decided to attach synI to synIII. Through the investigation of three-dimensional structures of fusion chromosome strains, unexpected loops and twisted structures were formed in chrIII-I and chrIX-III-I fusion chromosomes, which depend on silencing protein Sir3. These results suggest a previously unappreciated 3D interaction between *HMR* and the adjacent telomere. We used these fusion chromosomes to show that axial element Red1 binding in meiosis is not strictly chromosome size dependent even though Red1 binding is enriched on the three smallest chromosomes in wild-type yeast, and we discovered an unexpected role for centromeres in Red1 binding patterns.

## INTRODUCTION

Budding yeast has 16 linear chromosomes, with chromosome sizes ranging from 0.23 Mb to 1.5 Mb. The relatively small genome (12Mb), a relatively high chromosome number (n=16) and its unique life cycle (mitotic proliferation in both haploid and diploid states) make *Saccharomyces cerevisiae* a convenient model organism for a global synthetic eukaryotic genome project. The Synthetic Yeast Genome Project (termed Sc2.0) was launched in 2011 with the completion of the first synthetic chromosome arms, chrIXR and semi-chrVIL(Dymond et al., 2011). Since 2011, six full-length synthetic yeast chromosomes have been successfully synthesized (Annaluru et al., 2014, Mitchell et al., 2017, Shen et al., 2017, Wu et al., 2017, Xie et al., 2017, Zhang et al., 2017). Through building those synthetic yeast chromosomes, new insights into gene expression regulation and genome structure were revealed, as well additional evidence of the flexibility of the yeast genome to adapt to genetic changes.

Interestingly, even though there are thousands of designer changes in synthetic yeast chromosomes, the 3D structures of synthetic chromosomes do not change much, except in the case of relocating ribosomal DNA (rDNA) arrays (Mercy et al., 2017). The relocation of rDNA from its native location – chrXIIR (a long arm) to a short arm (chrIIIR) splits chrIII right arm into two non-interacting regions and imposes some new restrains to the nuclear genome. However, those synthetic yeast grow fine even with changes in 3D structure after rDNA relocation. It indicates that yeast could tolerate big changes in nuclear 3D architecture. In other cases, yeast 3D structure was probed to be related to its function. For example, distinct conformations of chromosome III observed in the two mating types may contribute to the donor preference phenomenon (Belton et al., 2015). Thus, the role of yeast 3D structure in transcriptome regulation and fitness remains of great interest.

During meiosis, small chromosomes tend to have higher rates of meiotic crossover, which is termed “chromosome size dependent control” (Kaback et al., 1992). This phenomenon was suggested to be regulated by crossover interference (Kaback et al., 1999). However, not every chromosomal segment follows this rule (Turney et al., 2004, Kaback et al., 1999). It has remained unclear whether these effects reflect some sort of direct or indirect measurement of chromosome size or whether this effect partially manifests through specific *cis*-acting elements. In this study, we complete the synthesis of synI, which is the smallest chromosome in wild type yeast. We also generate a series of wild-type fusion chromosome strains to probe the impact of chromosome structure on three-dimensional genome structure as well as meiotic function.

## RESULTS

### The design of synI

Compared to other Sc2.0 chromosomes, synthetic chromosome I (synI) has many unique features. First, synI is dramatically shorter than its native counterpart (180,554bp vs 230,208bp), yielding a relative length reduction of 21.6%. This contrasts other synthetic chromosomes that are on average only 8% shorter than wild type (Richardson et al., 2017). This is due in large part to the removal of two ∼10 kb subtelomere repeats, called W’ sequences, which lack coding sequences. It was previously suggested that W’ repeats might function as “filler” DNA to increase the size and stability of this smallest yeast chromosome (Bussey et al., 1995). Since chromosome I is the smallest chromosome in *S. cerevisiae*, more than 6x shorter than the largest chromosome (IV), we were concerned that further shortening it would negatively affect its stability (Bussey et al., 1995, Murray et al., 1986). Thus, a second distinct design feature of synI is that its design specifies attachment to another synthetic chromosome to help ensure its stability. To this end, synI does not encode a centromere and only the “unattached” right end is designed with a telomere. A second benefit of appending synI to another chromosome is that it balances the Sc2.0 karyotype at n=16, which would otherwise be increased by 1 due to the planned introduction of a supernumerary tRNA neochromosome (Richardson et al., 2017).

Besides these unique properties, synI design adhered to the general principles laid out for Sc2.0 chromosomes (Dymond et al., 2011, Richardson et al., 2017) (Fig. 1). Specifically, 4 tRNAs, all introns, and all retro-transposable elements were deleted. Further, 3535 bp were recoded for PCRtags, enabling differentiation of synthetic and wild-type DNA in a PCR-based assay (Dymond et al., 2011) and 210 bp were recoded to create or eliminate restriction enzyme cut sites, used for chromosomal assembly. 19 TAG stop codons were swapped to TAA, to free up one stop codon for future amino acids expansion. 62 loxPsym sites were added 3bp downstream of non-essential gene stop codons along with other major landmarks, such as telomeres and sites of tRNA and repeated DNA deletion sites.

**Figure 1.**
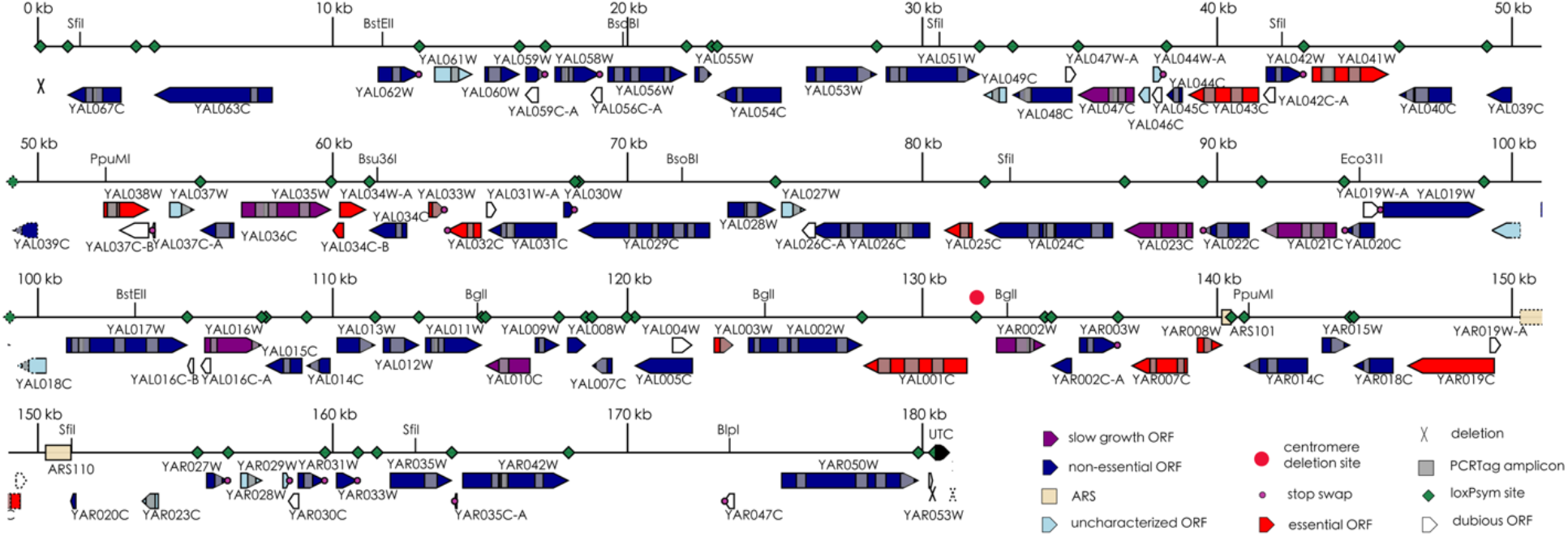
SynI design. SynI (180,554 base pair) encodes all Sc2.0 design features, with a relative length reduction of 21.6% compared to wild type chr1. A synthetic Universe Telomere Cap replaces the wild type telomere, and the large deletions at subtelomeres are marked by “X”. *CEN1* was removed allowing synI to be attached to another synthetic chromosome. All tRNAs and retrotransposable elements were deleted, as well as introns. 19 TAG stop codons were recoded to TAA and 62 loxPsym sites were added to the 3’ UTR of non-essential genes.

### Strategy to fuse chrI to three different chromosomes

The first major decision to be made prior to synI construction was the chromosome to which it should be attached. We did not know whether the altered 3D environment forced by such attachment would affect chrI performance, so we set out to fuse chrI to three chromosomes with distinct arm lengths: chrIIIR, chrIVR and chrIXR (Fig. 2a). chrIXR is the shortest arm in *S. cerevisiae* with a length of ∼84 kb, while chrIVR is the longest arm (excluding the variable-length rDNA locus on chromosome XII), at over 1 Mb. ChrIIIR has an intermediate length of ∼202kb. To evaluate the performance of these fusion chromosomes, we used wild type chromosomes.

**Figure 2.**
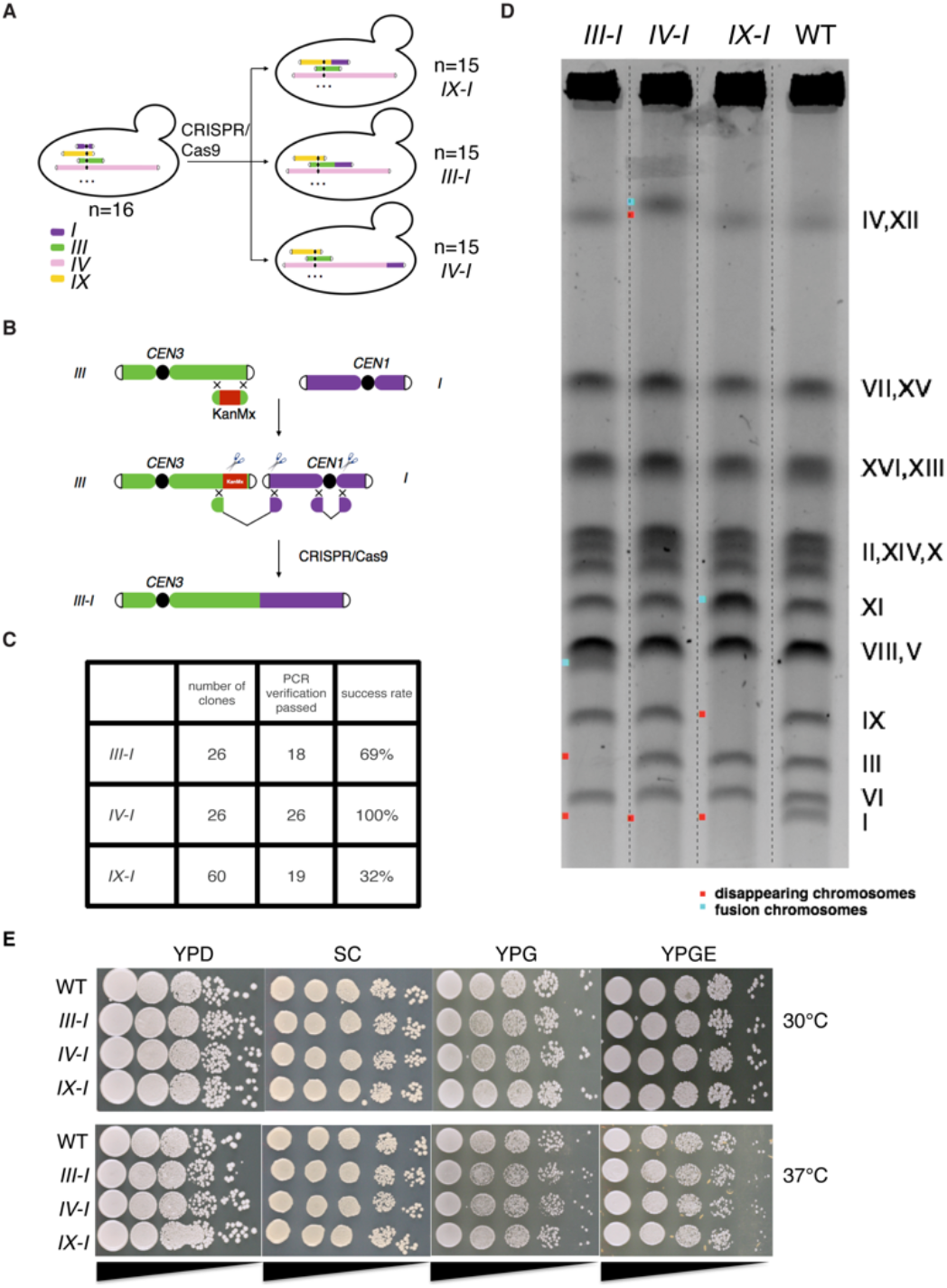
Construction and characterization of wild-type fusion chromosome strains. A, Schematic outlines the strategy used to construct chromosome I fusion strains. B, Schematic showing a CRISPR-Cas9 method deployed to fuse chrI to other chromosomes. C, The efficiency of the fusion chromosome method for chrI. D, Pulsed field gel electrophoresis, fusion chromosome strains are indicated. WT, wild-type strain Red dot = former location of wild-type chromosomes, blue dot = new fusion chromosomes. E, Serial dilution assays to evaluate fitness of fusion chromosome strains. YPD, yeast extract peptone dextrose. SC, synthetic complete medium. YPG, yeast extract peptone with 3% glycerol. YPGE, yeast extract peptone with 3% glycerol and 3% ethanol.

### Characterization of fusion chromosomes

To fuse chromosomes in *S. cerevisiae*, we used a previously developed variation of a CRISPR-Cas9 based method that allows us, in principle, to fuse chrI to any other chromosome in budding yeast (Fig. 2b) (Luo et al., 2018). For synI, this method was very robust, with success rates varying from 32% to 100% (Fig. 2c). Pulsed field gel electrophoresis was used to karyotype fusion chromosomes (Fig. 2d). ChrI and its fusion partners (chrIII, chrIV or chrIX) showed the expected, slower migrating bands (chrIII-chrI, chrIV-chrI or chrIX-chrI). To evaluate the growth fitness of the three fusion chromosome strains, serial dilution dot assays on different media and temperature conditions were performed (Fig. 2e). Under all the conditions tested, fusion chromosome strains grew as well as wild type, without any observed differences, consistent with previous observations (Luo et al., 2018).

### Assembly of synI

That all three fusion chromosome strains grew as well as wild type indicates that the attachment of chrI to different length chromosome arms does not affect the overall performance of chrI. For the purpose of moving ahead with synthesis and assembly of synI, we decided to attach chrI to synIII for two reasons. First, synIII was successfully assembled at that time, while synIX and synIV were still under construction. Second, synIII demonstrates high fitness, as tested under more than 20 conditions, including elevated temperatures, DNA replication stress, DNA damage stress and oxidative stress conditions (Annaluru et al., 2014). In just one out of 21 conditions tested (high sorbitol), synIII grew a little bit slower. To assemble synI, we started with a synIII-wt chrI fusion chromosome strain, containing the synIII centromere as the sole centromere, generated using the CRISPR-Cas9 method. This enabled subsequent use of the classic “Switching Auxotrophies Progressively for Integration (SwAP-In) method, allowing us to replace wild type chrI with synthetic DNA, in a series of 5 steps (Fig. 3a) (Richardson et al., 2017).

**Figure 3.**
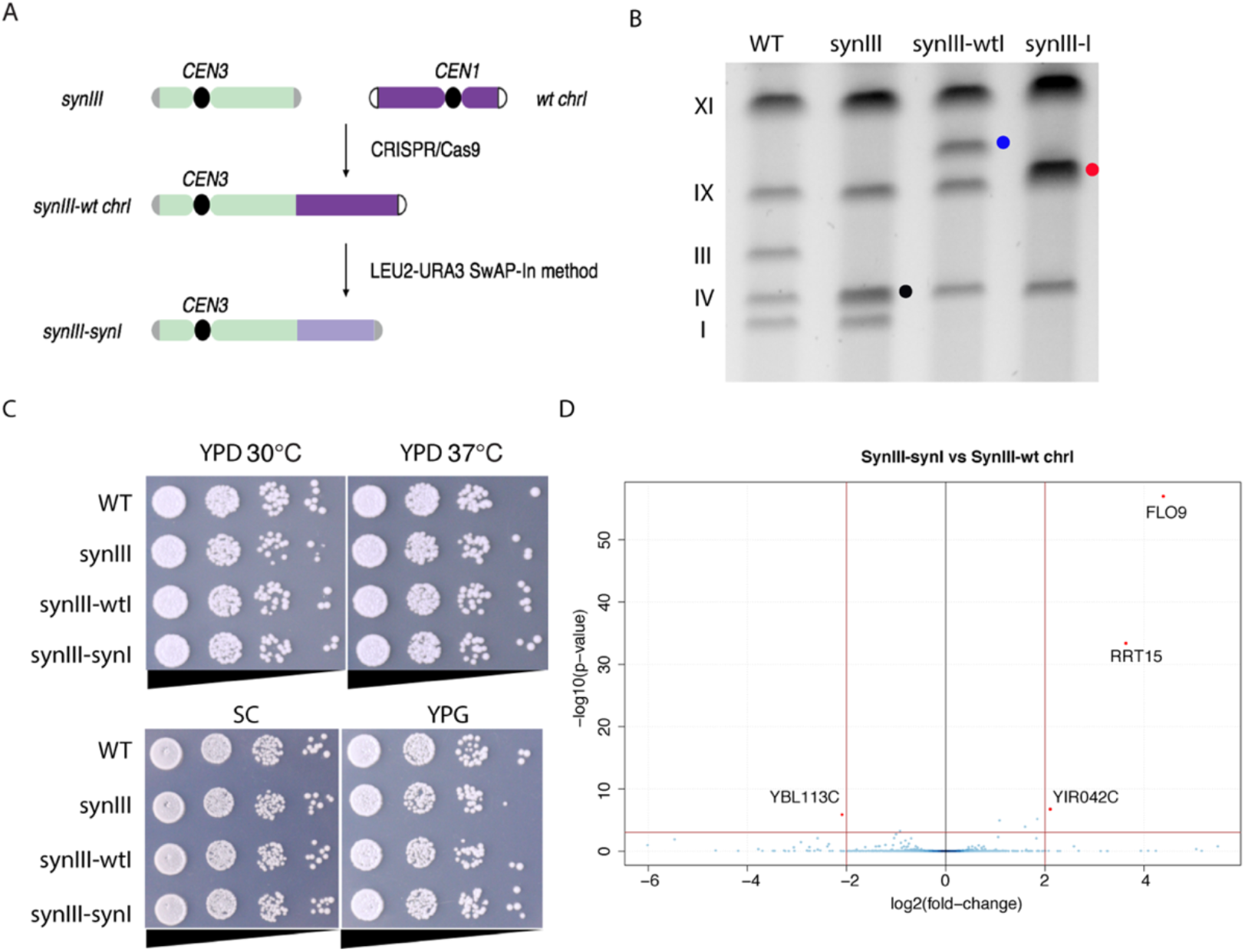
Construction and characterization of synthetic chromosome I (synI) A, Schematic showing the strategy to assemble synthetic chromosome I. B, Pulsed field gel electrophoresis result. SynIII (∼ 273 kb, marks by black dot) migrates faster than wild type chrIII (∼ 317 kb). The attachment of wild type chromosome I to synIII (∼ 496 kb) migrates slower, marked as a blue dot. Red dot indicates size of synIII-synI fusion chromosome (∼453 kb). C, Serial dilution growth assay. YPD, yeast extract peptone dextrose. SC, synthetic complete medium. YPG, yeast extract peptone with 3% glycerol. D, A volcano plot showing RNA seq data by comparing the transcriptome of synIII-synI and synIII-wt chrI strain. Red dots indicate genes whose expression is significantly different in the synIII-synI strain compared to synIII-wt chrI strain (p<10^−3^, log_2_|fold change| >2).

After attachment of native chrI to synthetic chromosome III, and five consecutive megachunk integrations, the ∼180 kb synthetic chromosome I was completed. PCRTags were used to detect the replacement of wild type genome by synthetic sequences (Supplemental Figs. 1 & 2). Pulsed field gel electrophoresis demonstrated that we successfully assembled fusion chromosome synIII-I (Fig. 3b). SynIII migrates faster than wild type chrIII due to a reduction in total length of ∼44 kb. In the synIII-wt chrI fusion strain, both chrI and synIII bands relocate to the predicted electrophoretic mobility of the synthetic fusion chromosome. The synIII-I chromosome (∼453 kb) co-migrates with chrIX (∼440 kb). During assembly, we identified a handful of synthetic features that caused apparent fitness defects and we elected to keep the wild-type sequence to preserve high fitness (Supplemental Table 1). The fusion chromosome strain, synIII-synI, grew without any defect compared to wild-type control strains, despite >20% reduction in chrI length and the removal of W’ repeats (Fig. 3c).

We performed whole genome sequencing to detect potential genome variation and RNA sequence (RNA seq) profiling to detect potential transcriptome changes. DNA sequencing of synIII-synI genome revealed a few SNPs and indels in synI compared to the designed genome (Supplemental Table 2). RNA seq revealed only four genes whose expression changed significantly. Among these, two are putative proteins with unknown function, one encodes Helicase-like protein within the telomeric Y’ element on chromosome 2 (*YBL113C*) and its expression was significantly reduced. Another, located on synI itself, encodes a lectin-like protein involved in flocculation (*YAL063C*-*FLO9*) (Fig.3c). *FLO9* is located on the left arm of chrI, near the chrIL telomere, which increased expression substantially. In the synI design, *FLO9* was recoded (using “REPEATSMASHER” (Richardson et al., 2017)) to facilitate proper assembly because native *FLO9* contains tandemly repeated peptide sequences (Verstrepen et al., 2005). Prior work has shown that telomere located genes are overexpressed when the telomere is fused to another chromosome (Luo et al., 2018, Shao et al., 2018) or the chromosome is circularized (Mitchell et al., 2017, Xie et al., 2017). However, it is theoretically possible that the extensive recoding done in *FLO9* could contribute to all or part of the increased expression.

### Chromosome fusion does not affect the overall 3D genome structure

To investigate potential changes of 3D nuclear architecture in wild type fusion chromosome strains. we exploited Hi-C to study three-dimensional organization of genomes (van Berkum et al., 2010). Wild-type yeast genome organization adopts a Rabl like configuration: a polarized array with 16 centromeres held together by tethering to a spindle pole body and telomeres within the opposing hemisphere (Duan et al., 2010). In this configuration, chromosome arms with similar lengths tend to interact more extensively (Therizols et al., 2010). Fusing chrI to another chromosome does not change the overall Rabl configuration (Fig. 4a). As expected, the attachment of chrI to another chromosome significantly increased both the local and the global (i.e. non-adjacent) interaction between chrI and the attached chromosome, indicated by higher contact frequency in 2D maps (Fig. 4a). An increased frequency of inter-chromosome interactions with chromosome arms of similar lengths to the “fusion arm” was also noted. In the most dramatic case, namely the chrI and chIV fusion, chrI interacted much more with long arms rather than short arms, revealing a completely different trend from the one in wild-type yeast (Fig. 4b).

**Figure 4.**
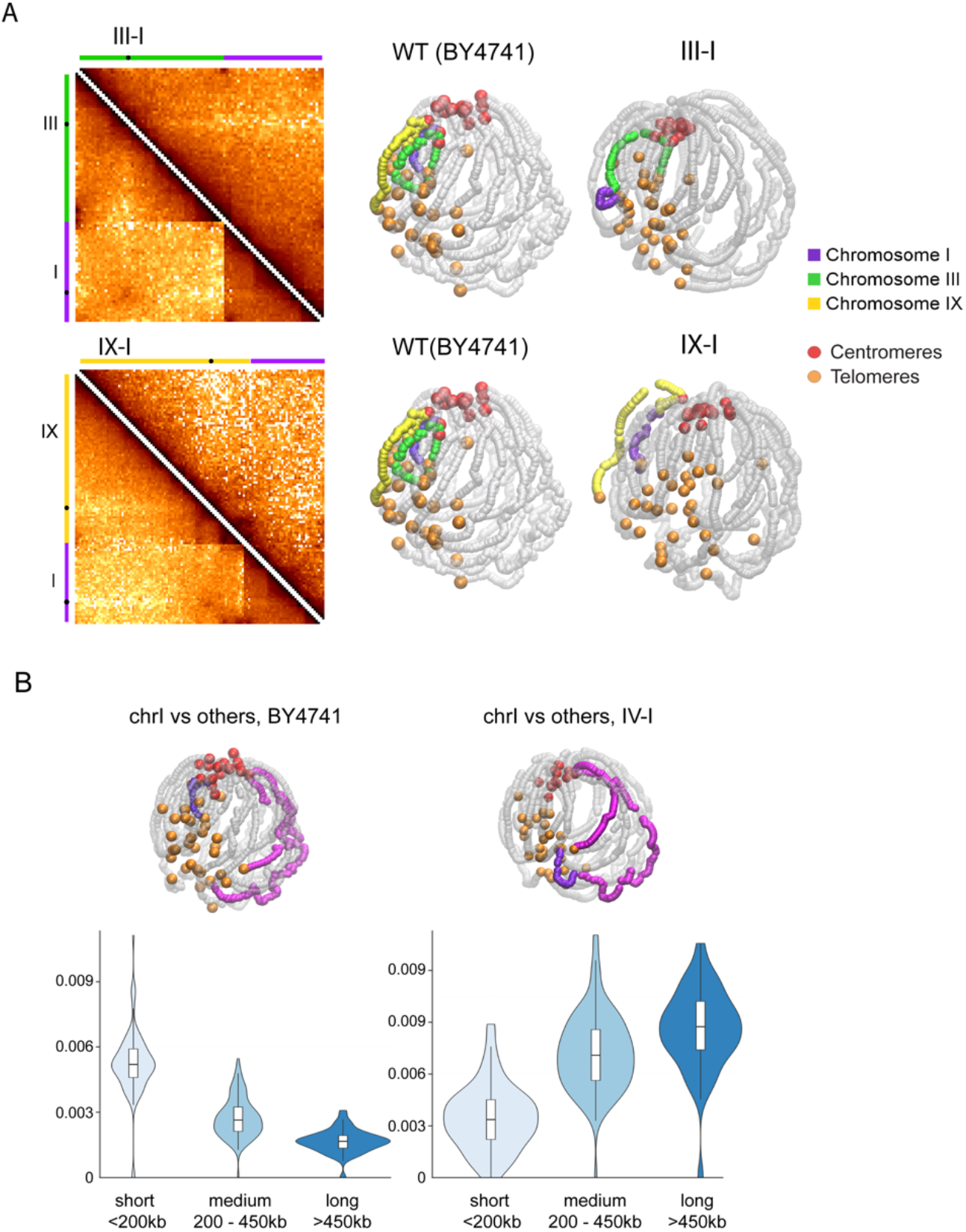
3D genome organization of wild type fusion chromosome strains. A, Comparisons of fusion chromosome strains and wild type strains normalized contact maps (5 kb bin) and 3D representations inferred from Hi-C contact maps. B, Violin plots showing the contact frequency of chrI to short (< 200 kb), medium (200 - 400 kb) or long (> 400 kb) chromosomal arms, for the wild type strain (BY4741) and for the strain with chrI fused to the long arm of chrIV. 3D representations are shown in the top of violin plots for visualization comparison; chrI is represented in purple and chrIV in pink, centromeres are colored in red and telomeres in orange.

### The chrIX-III-I fusion chromosome displays a Sir3-dependent complex twisted structure

In comparison to the wild type genome, where chrI stretches out in the typical Rabl configuration, in the chrIII-I fusion we observed an unexpected loop forming between the chrIR telomere and the fusion point, proximal to the *HMR* locus on chromosome III (Fig. 4a, chrIII-I strain). We did not see this loop in chrIX-I fusion chromosome strain, suggesting that cis-sequences on chrIII are required for the formation of this structure. *HML* and *HMR* are silent mating type cassettes, located near the left and right ends of chrIII, respectively, that are silenced by binding of the SIR protein complex (Rusche et al., 2003). In the chrIX-III-I strain, which was generated by fusing chrIXR to the left arm of chrIII-I, the chromosomal fusion points represent positions at which *HML* and *HMR* interact (Fig. 5a). The 3D representation suggests that the chrIXR sequences cross between chrIR and *HMR* sequences and are extruded, forming a complex twisted structure. This interaction pattern could be explained by a strong *HML-HMR* interaction leading to a geometric block of the chrI telomere loop. Interestingly, these novel twisted/looped structures were absent from *sir3Δ* variant of the chrIX-III-I strain, consistent with a dependence on the Sir protein silencing complex in the formation and/or maintenance of this 3D structure. The comparison map between chrIX-III-I strains with and without Sir3 revealed that the interactions between chrIX and chrI or chrIII were much stronger in the presence of Sir3 (Figs. 5b, 5c).

**Figure 5.**
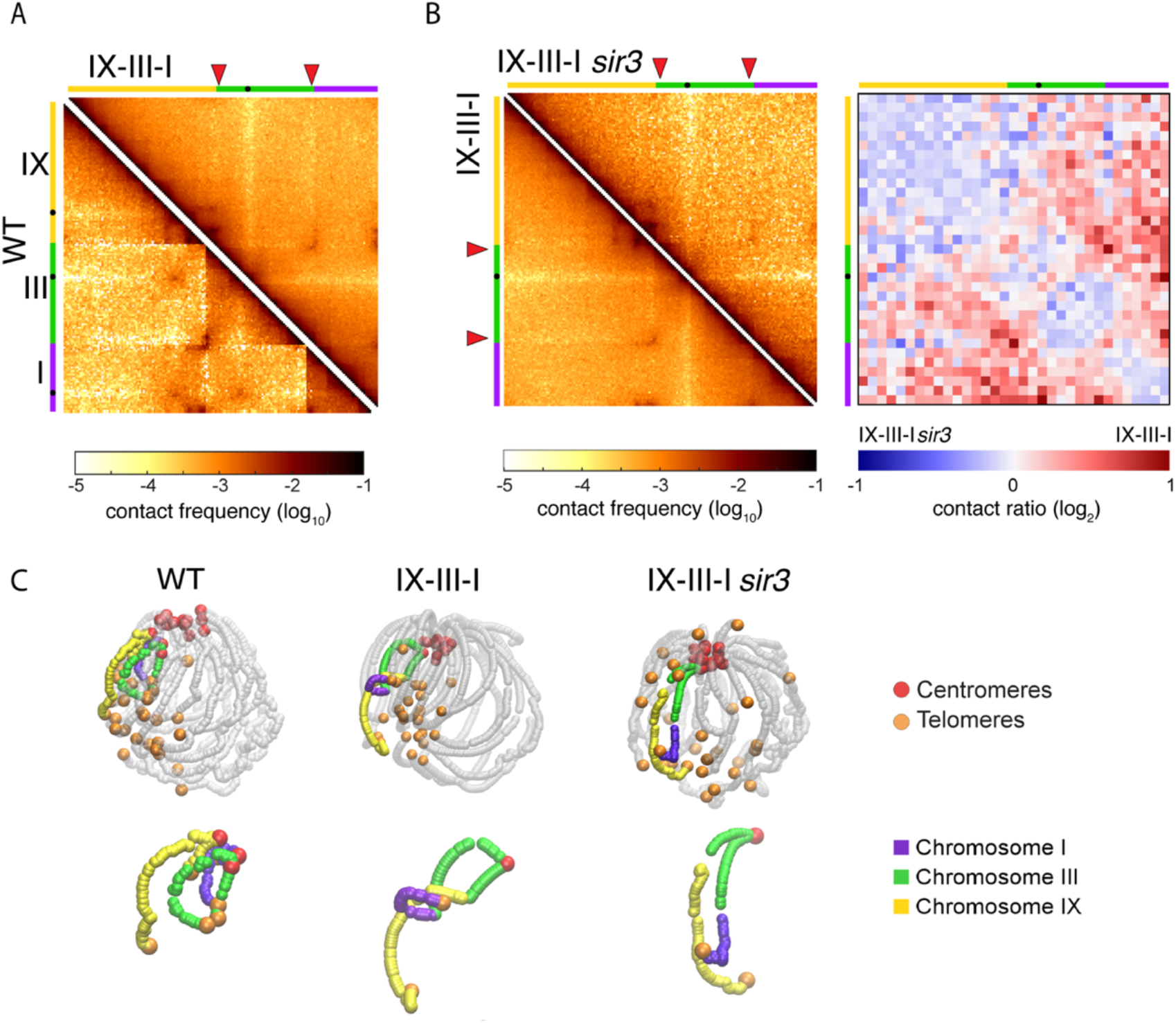
Twisted structure formed in chrIX-III-I is Sir3 dependent. A, Comparisons of normalized contact maps of wild type strains and chrIX-III-I strains. Red arrows point at HML and HMR loci. B, Comparisons of normalized contact maps of chrIX-III-I strains with or without Sir3 protein. Left panel: normalized contact maps. Right panel: ratio between two contact maps (50 kb bin). Blue contacts are stronger in the chrIX-III-I *sir3* strain, red contacts are stronger in the chrIX-III-I strain. C, 3D representations inferred from the contact maps using Shrek 3D (Lesne et al., 2014). The blank spaces in the junctions on chrIX-III-I *sir3* strain reflects the lower mappability of subtelomeric sequences, which were later excluded from subsequent analyses.

### Preferential Red1 binding to small chromosomes is linked to *cis*-acting elements

The fusion chromosomes also provided an opportunity to investigate effects of chromosome size on meiosis. Small chromosomes in many sexually reproducing organisms, including humans, exhibit elevated rates of recombination, which is thought to assist in their proper segregation during meiosis I (Kaback et al., 1992, Kaback, 1996). In yeast, this chromosome size preference is reflected in a meiotic enrichment of the recombination-promoting proteins Red1 and Hop1 on small chromosomes (chrI, III and VI) (Sun et al., 2015, Panizza et al., 2011). It is unclear, however, whether this preference is the result of chromosome-encoded DNA elements or mechanisms that respond to chromosome size. To investigate this question, we analyzed Red1 binding on wild-type and fusion chromosomes by ChIP-seq. This experiment was performed in hybrid BY4741/SK1 strains, which bypass the poor sporulation efficiency of BY4741 (Deutschbauer and Davis, 2005). The presence of the same SK1 genome in all hybrids also provides an internal control of ChIP-seq efficiency, allowing normalization between samples.

The ChIP-seq data show that the chrIV-I strain in which chrI is attached to chrIV (and centromere IV is deleted) did not change total Red1 binding to chrI sequences despite the massive size increase (Fig. 6c). Total Red1 binding on chrIII sequences also did not change in the chrIX-III-I fusion chromosome strain, with intact centromere III. These data suggest that biased enrichment of Red1 on small chromosomes is not linked to chromosome size and implies a role for *cis*-acting elements.

**Figure 6.**
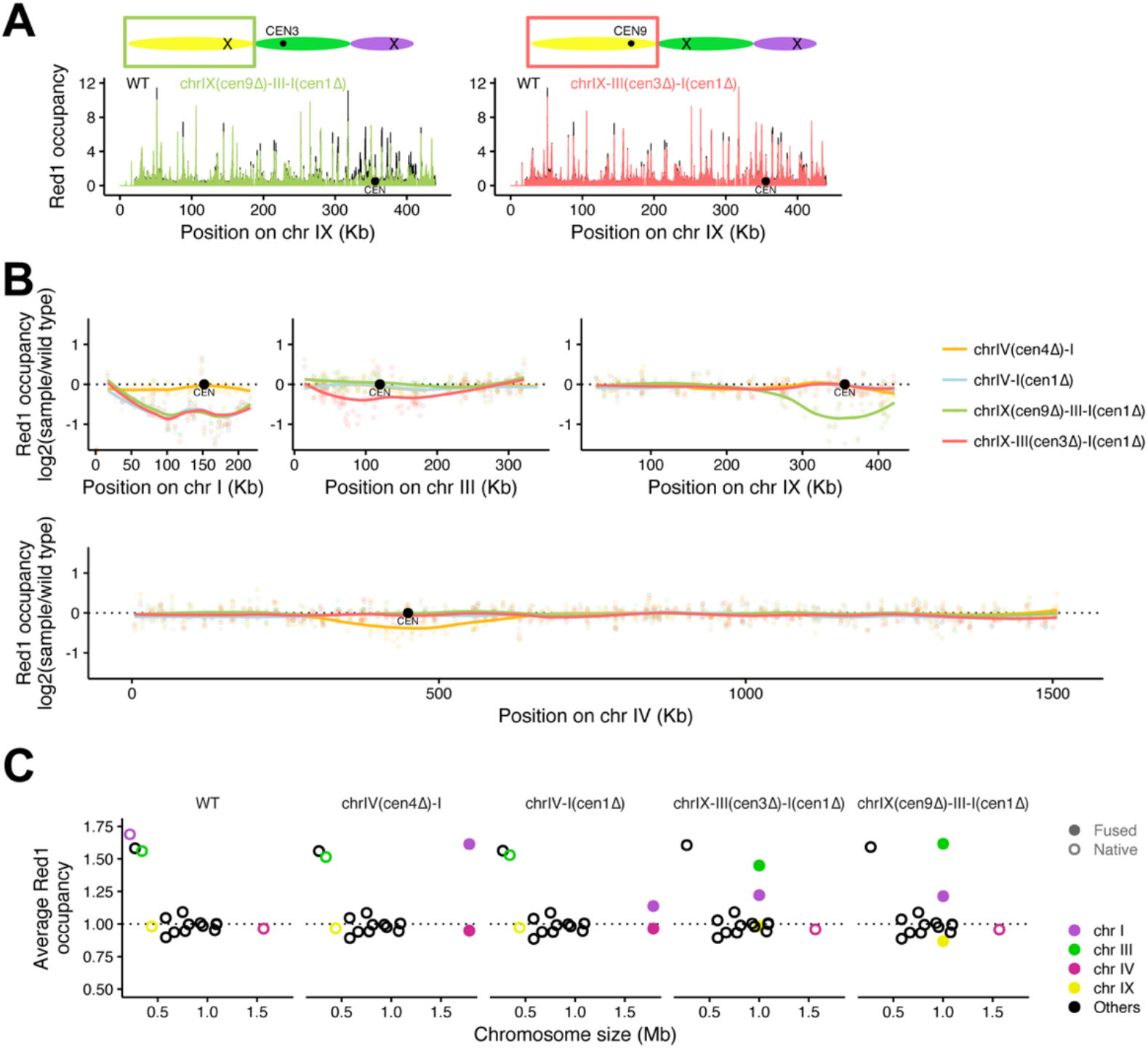
Meiotic axis protein Red1 occupancy on wild type fusion chromosome strains. Red1 occupancy is represented by the input and sequencing depth-normalized fragment pileup (fragment pileup per million reads), further normalized to mean Red1 occupancy on SK1 chromosomes. A, Red1 occupancy along S288C chromosome IX for each fusion chromosome strain overlaid on the wild type occupancy. B, Mean signal in Red1 peaks along fusion chromosomes as log2-transformed ratios between each strain and the wild type. Points represent mean peak signal and lines represent local regression (loess normalized). C, Red1 occupancy versus chromosome length in the S288C background.

### Red1 binding is affected locally by loss of centromere activity

To identify such *cis*-acting elements, we analyzed Red1 binding along chromosomes. The position of Red1 binding sites was unchanged between wild type and fusion chromosomes (Fig.6a). However, we noted a highly reproducible local drop in Red1 signal intensity whenever a centromere was deleted in the construction of a fusion chromosome (Figs. 6a, Supplemental Fig.3). For example, when *CEN9* was retained in the construction of chrIX-III-I, we observed a drop in overall Red1 signal around the inactivated *CEN3* and deleted *CEN1*. Conversely, when *CEN3* was retained in the construction of chrIX-III-I, signal dropped around the deleted *CEN1* and *CEN9*. This effect is also seen for *CEN4* (Figs. 6b, Supplemental Fig.3) and thus also occurs at centromeres of large chromosomes.

Plotting signal within Red1 peaks as a fraction of wild type revealed a decrease in Red1 signal across a region of approximately 200 kb centered on the deleted centromeres (Fig. 6b). The range of this effect is extraordinary given that in some cases only a couple of nucleotides were mutated to inactivate a centromere (Supplemental Table 4). These data suggest that the drop in Red1 signal is a consequence of losing centromere activity rather than alterations of base composition or other larger scale consequences of sequence deletion. The large sequence range over which this effect is seen has a profound impact on the enrichment of Red1 on small chromosomes, which, in the case of chrI, are almost completely encompassed by this effect. Accordingly, when *CEN1* is deleted in the context of a fusion chromosome, average enrichment of Red1 on chrI sequences drops near to the genome average (Fig. 6c). By contrast, deletion of *CEN4* had only mild effects on the average Red1 enrichment on the sequences of this much larger chromosome. Thus, centromere inactivation largely eliminates the biased enrichment of Red1 on small chromosomes during meiosis.

## DISCUSSION

As a consequence of building synthetic chromosome one, synI, as part of the synthetic yeast genome project, Sc2.0, we have explored in detail a set of fusion chromosome strains. We attached chrI to three separate chromosomes with arms of different lengths, dramatically changing the neighbors that chrI interacted with most frequently in *trans*. In the extreme case, when attached to the long arm of chrIV, chrI interacted strongly with all long chromosome arms and much less with small arms. This is completely opposite of what is seen with the native configuration of chrI. This finding provides direct evidence for a previous study related to DNA repair and genomic environment: the relocation of the same fragment to different length chromosome arms changes the environment of the fragment, leading to differential efficiencies of homology searching and DNA repair (Agmon et al., 2013).

The yeast also grew very well even with a significantly reorganized 3-dimensional genome structure. These findings reveal extreme tolerance of the yeast genome with respect to 3D positioning inside the nucleus. Previous work supporting this idea is provided by a study in which the rDNA cluster was relocated from chrXII to the much shorter chrIII (Zhang et al., 2017). The right arm of chrIII was split into two, no-longer interacting segments due to the insertion of the 1-2 Mb rDNA array. No growth fitness defects were detected in that yeast strain either, despite a major rearrangement of multiple yeast chromosomes to accommodate the ectopically located rDNA (Mercy et al, 2017). A recent study also showed that yeast grow well even with dramatic rearrangements of yeast 3D genome structure by fusing 16 chromosomes into 1 or 2 giant chromosomes (Shao et al., 2018, Luo et al., 2018). Together, these data reinforce the fact that the budding yeast is extremely plastic, and can tolerate surprisingly substantial changes in 3D genome structure without obvious ill effects.

When we performed Hi-C analysis to study the 3D structure of the three fusion chromosome strains in mitotic cells, we observed an unexpected loop form between *HMR* and the chrIR telomere of the chrIII-I fusion chromosome. This loop further constrains the chrIX-III-I fusion chromosome into a more twisted configuration. Specific long-range interactions between *HML* and *HMR* are known to depend on SIR family silencer proteins (Miele et al., 2009). A previous study revealed a loss of specific long-range interactions between the two silent mating type loci in synthetic chromosome III strains when *HML* and *HMR* were removed by design (Mercy et al., 2017). Telomere silencing also requires the binding of Sir proteins. Confocal microscopic images show that in normal yeast strains telomeres are clustered into 6-8 foci at the nuclear periphery, sometimes overlapping with *HML* and/or *HMR* (Taddei and Gasser, 2012). Therefore, we hypothesize that the unexpected loop and twisted structures observed represent previously unknown *HMR, HML* and telomere interactions that are either held together or maintained at the nuclear periphery, by silencing protein complexes. Consistent with this, when we depleted Sir3 in the chrIX-III-I strain, the twisted structure was no longer observed.

In meiosis, chromosomes are packaged in a distinct manner to assure proper disjunction of homologs and the formation of double-strand breaks and subsequently, chiasmata. Specific proteins mediate the formation of meiosis I-specific chromatin. Meiotic recombination rates, however, were previously shown to be chromosome size dependent (Kaback et al., 1992, Kaback, 1996). The recombination rates between the same loci increased when chrI was split into two even smaller segments, while they decreased when a longer chrI was constructed via a reciprocal translocation of the chrII left arm. However, as not all chromosomal segments follow this pattern (Turney et al., 2004, Kaback et al., 1999), it has remained unclear whether these effects reflect a measurement of chromosome size or the action of *cis*-acting elements. Previous studies showed that the binding of Red1 protein is enriched on three small chromosomes (Sun et al., 2015), correlating positively with a higher frequency of double-strand breaks (Blitzblau et al., 2007, Pan et al., 2011) and higher meiotic crossover rates on small chromosomes (Kaback et al., 1992). Our analysis of fusion chromosomes shows that the relative enrichment of Red1 on small chromosomes (chrI, chrIII) is retained when these sequences are placed into the context of much larger chromosomes, strongly suggesting a role for *cis*-acting elements in elevating Red1 binding levels on small chromosomes.

Intriguingly, deletion or inactivation of centromeres on small chromosomes led to a profound reduction in Red1 binding on fusion chromosomes. Decreased Red1 binding extended approximately 100 kb to either side of the inactivated centromere. Although all tested centromeres show this effect, the long-range nature of the Red1 binding disproportionately affects small chromosomes. This provides an interesting paradigm for how a genetic element present on all chromosomes can nevertheless have a much more pronounced effect on small chromosomes. Centromere deletion might also delay the early pairing of these sequences (Tsubouchi et al., 2008) which could effect overall recombination rates. Such an effect might be quantitatively confounded in either direction by the fact that only examined cells were arrested at a single time point.

The source of the observed decrease in Red1 levels, however, remains unclear. Red1 enrichment within 100kb of centromeres is not elevated over the genome average on wild-type chromosomes (Sun et al., 2015), indicating that centromeres do not simply cause increased deposition of Red1 protein. We note that in our experiments, the centromere-deleted regions of fusion chromosomes were present in diploids that also harbored wild-type chromosomes, and here were thus in a hemizygous state. As homologous chromosomal sequences become intimately paired in the context of meiotic prophase (Zickler and Kleckner, 2015), it is possible that the presence of an active centromere on one homolog can influence the Red1 enrichment signal observed on a paired homolog lacking a centromere. Alternatively, the changes in Red1 signal may reflect differences in chromosome organization mediated by centromere-proximity that may be captured by the crosslinking prior to ChIP-seq analysis. Additional analyses will be necessary to distinguish between these possibilities.

## ACKNOWLEDGEMENTS

We thank the NYU Langone Health Genome Technology Center, especially A. Heguy, G. Westby and P. Zappile, for deep sequencing libraries, and Z. Kuang for advice on Bioinformatics analysis. This work was supported by NSF grant MCB-1616111 to J.D.B., NSF grant MCB-1445545 to J.S.B. and funding from the National Institutes of Health (1R01GM111715) and the March of Dimes Foundation (6-FY6-208) to A.H.

## AUTHOR CONTRIBUTIONS

J.L., and J.D.B. conceived the project. L.A.M and J.D.B designed synI. J.L., R.K. A.H. and J.D.B. designed and analyzed the experiments. J.L., G.M. (in the lab of R.K.), L.A.V.S. (in the lab of A.H.), W.Z., N.A., J.Y.A., G.A., A.D., H.B., Y.C., M.C., O.F., R.V.G., S.K., S.K., H.S.L, J.I., L.M., A.M. S.M., N.M., P.N., M.N., A.R., A.T., T.W., N.W., T.Z., V.Z. and K.Z. performed the experiments. G.S. and K.Y. (in the lab of J.S.B.) performed chromosome segmentation and carried out computational analyses. X.S. (in the lab of J.D.B.), L.A.V.S. (in the lab of A.H.), G.M (in the lab of R.K.) and J.M. performed computational analyses. J.L and J.D.B wrote the manuscript, and all authors contributed to its editing.

## STAR METHODS

### Strains

All strains generated in this study are listed in Supplemental Table 3. For wild type fusion chromosome strains, they are all derived from BY4741 (*MATa his3Δ0 leu2Δ0 met15Δ0 ura3Δ0)* by CRISPR-Cas9 editing. For synI strain construction, the starting strain is synIII (YLM422: *MATα his3Δ1 leu2Δ0 lys2Δ0 ura3Δ0 SYN3-272123bp HO::synSUP61*) (Annaluru et al., 2014). For Red1 chip-seq experiments, hybrid diploids were generated by mating fusion chromosome strains (*MATa*) to SK1 strain (*MATα ho::LYS2, lys2, ura3, leu2::hisG, his3::hisG, trp1::hisG*).

### CRISPR-Cas9 method to fuse chromosomes

A KanMX cassette was first inserted into the subtelomere region of a chromosome arm to which chrI was to be fused. Then we used a previously developed CRISPR-Cas9 (Luo et al., 2018) to generate cuts near the chrIL telomere, *CEN1* and inside the *KanMX* coding sequence, and provided two donors: one donor with ∼400bp homology sequences to the chrIL subtelomere and the chromosome arm to be fused, and another donor carrying ∼400bp homology sequences flanking *CEN1*. The primers that we used to amplify 400 bp homology sequences are provided in Supplemental Table 5. The 20nt gRNA of KanMX is 5’-CTTCGTACGCTGCAGGTCGA-3’, and the 20nt gRNA of *CEN1* is 5’-AAGAAAGTTATATGTGTGAC-3’. Chr1L-gRNA sequence is 5’-CTCAATGTACGCGCCAGGCA-3’. Two telomeres and *CEN1* were deleted in each fusion experiment (Luo et al., 2018). We first screened colonies for “winner” candidates by replica plating to YPD+G418 to select colonies sensitive to G418, followed by PCR verifications as previously described (Luo et al., 2018). Note, for chrIX-I strain, it was generated by two steps yeast transformation rather than using CRISPR-Cas9. First step, we inserted a fragment carrying ∼400bp homology sequence to left arm of chrI and a *URA3* selection marker into chrIX right arm near telomeres by homology recombination. Second step, we transformed cells with a fragment containing a *HIS3* marker flanking by ∼400bp homology sequences around *CEN1*. Then we selected for cells that grow on SC–His plate and PCR verified the fusion of two chromosomes.

### Whole genome sequencing and RNA seq

The libraries preparation of whole genome sequencing and RNA seq, as well as Data analysis was performed as previously described (Luo et al., 2018), except that the libraries were sequenced as 150 bp single-end reads on an Illumina NextSeq 500. The thresholds we used for differentially expressed genes are p-value < 10^−3^ and log 2|fold change| > 2.

### Hi-C libraries

Hi-C libraries were generated using the DpnII restriction enzyme and following the same protocol as the one described (Lazar-Stefanita et al., 2017). The resulting libraries were used as template and processed for 2×75 bp pair-end sequencing on an Illumina NextSeq500 according to manufacturer’s instructions.

### Generation and normalization of contact maps

Illumina sequencing data were processed as described in Lazar-Stefanita et al. 2017. First, PCR duplicates were removed. Second, each read of a pair was aligned independently with Bowtie 2.1.0 (mode: --very-sensitive --rdg 500,3 --rfg 500,3) and using an iterative procedure as described (Imakaev et al., 2012) against the genome of the corresponding fusion strain. Third, unwanted molecules (*i.e.* corresponding to loops, non-digested fragments, etc.; for details see (Cournac et al., 2016, Cournac et al., 2012)) were discarded. Finally, the remaining valid reads were binned into units of single restriction fragments, then successive fragments were assigned to fixed size bins of 5 kb. Contact maps were then normalized using the sequential component normalization procedure (SCN)(Cournac et al., 2012). The contact ratios were computed between normalized maps binned at 50 kb.

### 3D representation inferred from contact maps

The 3D representations of the contact maps were generated using ShRec3D (Lesne et al., 2014) on the normalized contact maps filtered for low signal bins. Each element of the normalized contact maps was inverted to compute sparse distance matrices. These matrices were then completed using the shortest path algorithm. 3D coordinates were then determined by multidimensional scaling using the Sammon mapping (Morlot et al., 2016). All 3D structures were rendered using VMD (Humphrey et al., 1996).

### Synchronous meiosis and chromatin immunoprecipitation (ChIP)

To induce synchronous meiosis, strains were pre-inoculated at OD_600_ = 0.3 in BYTA medium (1% yeast extract, 2% tryptone, 1% potassium acetate, 50 mM potassium phthalate) and grown for 16.5 h at 30°C. Cells were then washed twice with water, resuspended at OD_600_ = 2.0 in SPO medium (0.3% potassium acetate, pH 7.0) and incubated at 30°C. After 3 h incubation, 25-ml samples were harvested and fixed for 30 min. in 1% formaldehyde. The formaldehyde was quenched with 125 mM glycine and samples were further processed as described previously (Blitzblau and Hochwagen, 2013). One tenth of the cell lysate was removed as an input sample and the remainder was immunoprecipitated for 16 h at 4°C with 2 μL of anti-Red1 serum (Lot#16440; kindly provided by N. Hollingsworth), followed by capture on GammaBind G sepharose beads (GE Healthcare Bio).

### Read alignment and peak calling

Sequencing reads were aligned to a concatenated *S. cerevisiae* SK1 and S288C strain hybrid genome using Bowtie v1.2.0 (Langmead et al., 2009). The hybrid genome was built by concatenating recently published assemblies of the SK1 and S288C genomes (Yue et al., 2017). Only reads aligning perfectly (no mismatches, flag “-v 0”) at a single position (flag “-m 1”) were considered. Reads that do not overlap a SNP between SK1 and S288C will align to both genomes, becoming multimappers which were discarded. Regions with enriched Red1 protein signal (Red1 peaks) were identified using MACS v2.1.1 (https://github.com/taoliu/MACS/; (Zhang et al., 2008)) with command “macs2 callpeak”. A “No shifting” model was built (flag “--nomodel”) and reads were extended towards the 3’ direction to a fixed fragment length of 200 (flag “--extsize 200”). Flag “--SPMR” was set, in order to generate fragment pileups per million reads. The mappable genome size was set to 24 Mb (flag “--gsize 2.4e7”) and default values were used for all other options. Input-corrected fragment pileups were generated with command “macs2 bdgcmp” and the fold enrichment method (flag “-m FE”). Raw sequencing data and fragment pileup files are deposited at GEO (GSE114731). A reviewer access token is created: epencuosbzydbgp. All code used to analyze the data is openly available online at https://github.com/hochwagenlab/Chr_fusion_hybrids.

