## supplemental figures 1 and 2 for "Synthetic chromosome fusion: effects on genome structure and function"

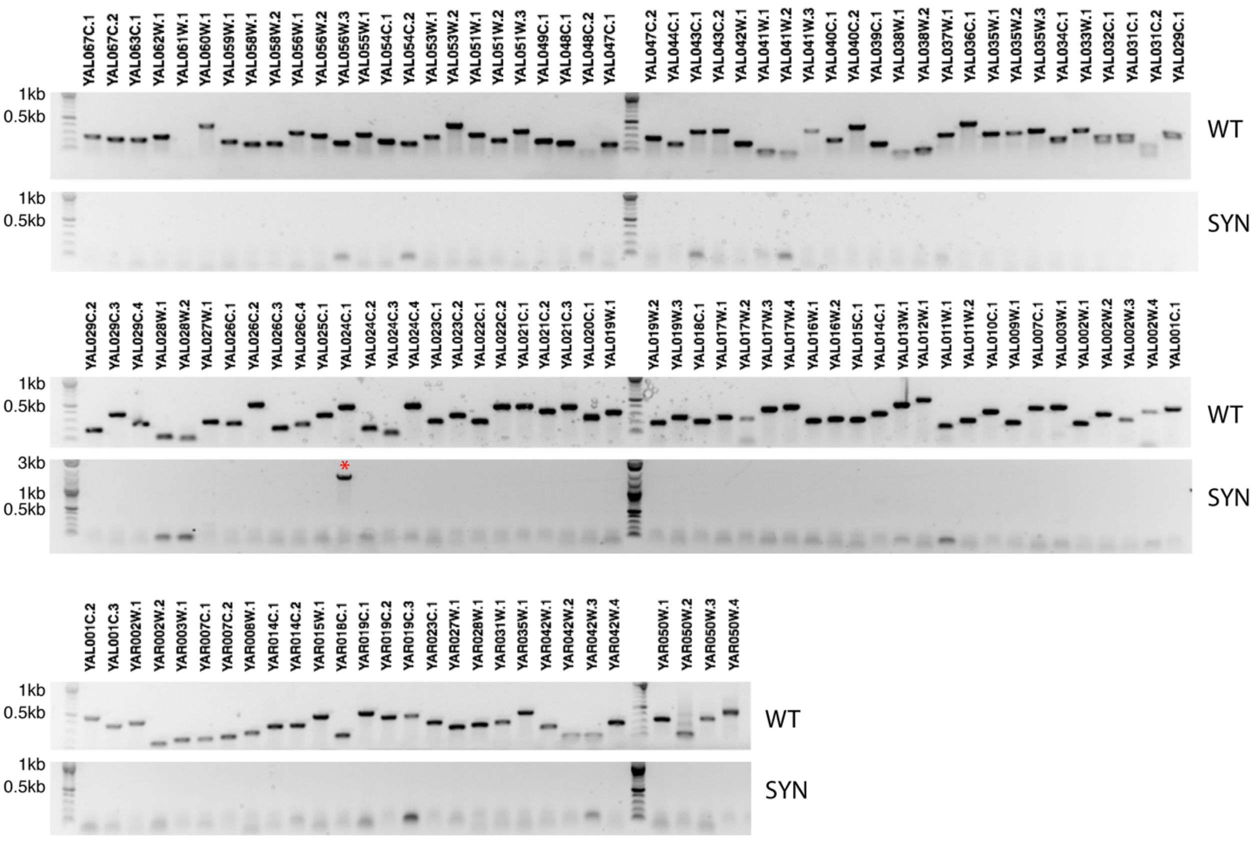


Supplemental Figure 1 PCRTags of wild type BY4741 genome

Wild type (WT) PCRTags and Synthetic (SYN) PCRTags were used to amplify the wild type genome. PCRTags amplicons are usually less than 500 bp. Only wild type PCRTags amplified amplicons from wild type genomic DNA except in one case where we observed a non-specific amplification with *YAL024C*.1 synthetic PCRtags with a size ~ 2 kb, which is indicated by *.

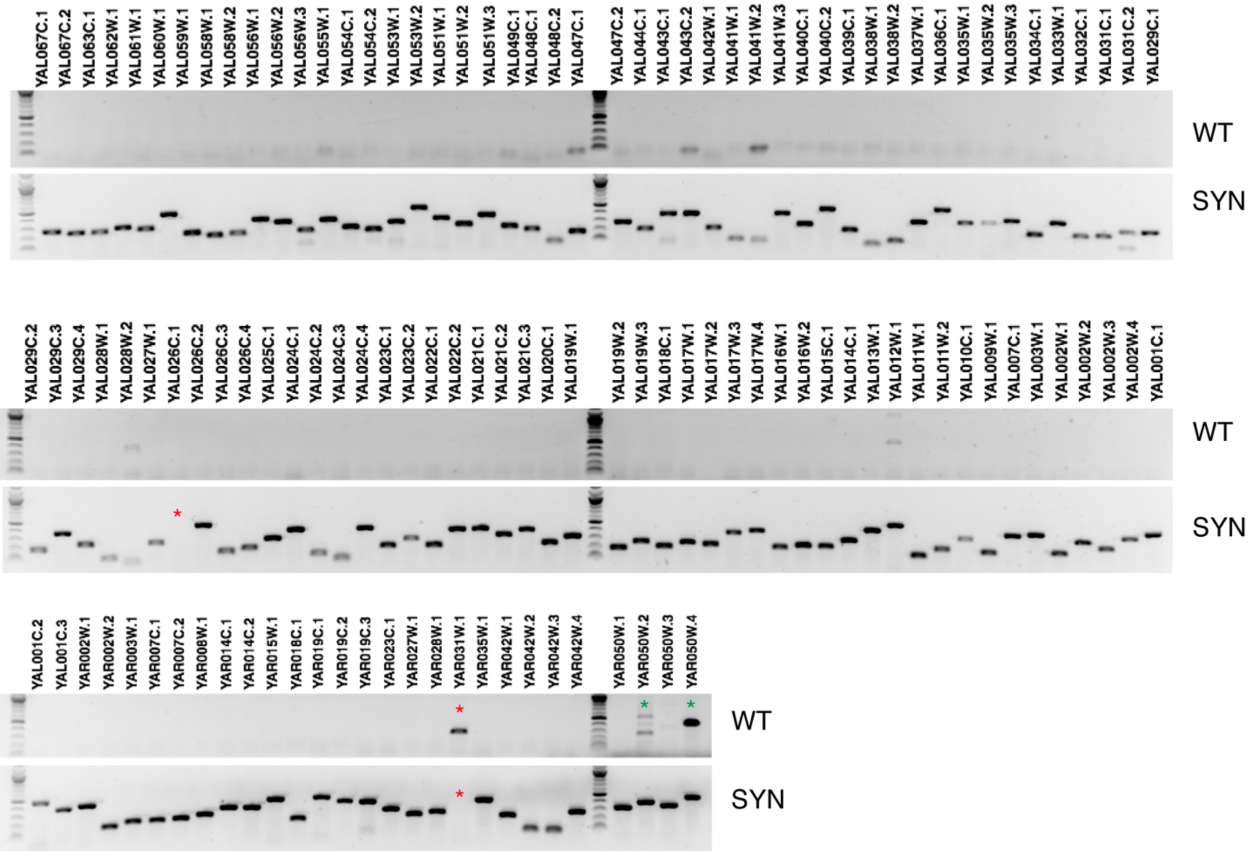


Supplemental Figure 2 PCRTags of *synI* genome

Wild type (WT) PCRTags and Synthetic (SYN) PCRTags were used to amplify the *synI* genome. PCRTag amplicons are usually less than 500 bp. There are PCRTags we intentionally kept as wild type because synthetic replacement caused growth defects, indicated by *. *FLO1*(*YAR050W*) has a paralog, *FLO5*, on *chrVIII*, to which *YAR050W.4* wild type PCRTags likely anneal, producing amplicons indicated by *.
